# Disease-specific differences in particulate matter handling drive pathogenic responses in human derived nasal epithelial cells

**DOI:** 10.64898/2026.05.20.726629

**Authors:** Yiran Li, Bangqiao Yin, Emily Eberhardt, Xinyi Deng, Rachael Bogle, Hannah L. Briggs, Christina E. Years, Jennifer Fox, Chad Brenner, Lam C. Tsoi, Johann E. Gudjonsson, Stuart Batterman, Mara Duncan, Marc B. Hershenson, Amarbir S. Gill

## Abstract

**Background:** Particulate matter (PM) exposure is associated with increased risk and exacerbation of chronic rhinosinusitis (CRS), yet underlying mechanisms remain poorly understood.

**Objective:** To define the epithelial mechanisms by which PM exposure promotes and exacerbates CRS, with a focus on epithelial remodeling, inflammation, barrier dysfunction, and cellular uptake of PM.

**Methods:** Human nasal epithelial cells obtained from ethmoid tissue of CRS (n = 5) and control donors (n = 4) were cultured at an air–liquid interface and exposed to PM. Single-cell RNA sequencing was performed to characterize PM-induced cellular and transcriptional changes. Protein expression, epithelial barrier integrity, cell death, and intracellular PM uptake were evaluated using biochemical, imaging, and ultrastructural approaches.

**Results:** Unsupervised clustering identified seven epithelial cell populations. Gene set analysis revealed baseline enrichment of inflammatory and keratinization pathways and reduced ciliogenesis in CRS compared with controls. Although PM induced inflammation and squamous differentiation in controls, the pathogenic responses were significantly amplified in CRS, including uniquely enhanced IL-1 signaling. Transcriptional changes were validated by ELISA, transepithelial electrical resistance, and immunofluorescence, demonstrating increased inflammation, epithelial barrier disruption, and cell death following PM exposure. Transmission electron microscopy revealed increased intracellular PM within membrane-bound organelles. Pre-treatment with an endocytosis inhibitor rescued PM-induced epithelial barrier dysfunction and inflammation.

**Conclusion:** CRS epithelium exhibits baseline dysfunction that may predispose it to environmental injury. PM exposure both induces CRS-like epithelial changes in controls and exacerbates disease-associated phenotypes.

**Key Messages:**

- Compared to controls, CRS nasal epithelium exhibits baseline inflammatory, keratinization, and ciliogenesis abnormalities.
- Particulate matter induces inflammation and squamous differentiation, while amplifying epithelial injury that is more robust in CRS epithelium compared to controls.
- Inhibition of dynamin-dependent endocytosis rescues PM-induced epithelial barrier leakiness and inflammation, implicating intracellular particulate matter uptake in disease pathogenesis.

**Capsule Summary:** Particulate matter induces CRS-like epithelial remodeling in controls and exacerbates inflammation and epithelial barrier dysfunction in CRS nasal epithelium, which can be rescued with endocytosis inhibition. This suggests a mechanistic link between baseline CRS vulnerability, intracellular uptake of particulate matter, and disease pathogenesis.

## Introduction

Chronic rhinosinusitis (CRS) is an inflammatory disorder of the upper airway affecting up to 12% of the U.S. population.^1–5^ Patients present with nasal congestion and drainage, smell loss, headache, and infections.^6^ Non-rhinogenic symptoms include fatigue, sleep and cognitive dysfunction, and depression/anxiety. Quality-of-life impairment is on par with coronary artery disease, end stage renal disease, and Parkinson’s disease.^7^ The overall annual economic burden of CRS in the U.S. is $22B.^8^ More than 250,000 endoscopic sinus surgeries are performed annually,^9, 10^ and 18% require revision.^11^

The etiology of CRS is thought to be multi-factorial, including contributions from occupational and environmental exposures.^6, 12^ For example, studies demonstrate a strong link between exposure to ambient particulate matter (PM) and the risk of developing CRS.^13–15^ Data also indicate increased disease severity and worse surgical outcomes among patients with CRS exposed to high levels of PM.^16, 17^ At the same time, wildfires, a major source of atmospheric PM,^18–20^ are increasing in frequency and intensity,^21–23^ and it is estimated that in the coming decade the wildfire burden on the healthcare system will be significant.^24^

PM can induce pleiotropic changes in human nasal epithelial cells (HNECs), including disruption of the epithelial barrier^25–29^ and induction of inflammation.^25, 30, 31^ PM has also been associated with impaired ciliogenesis and epithelial remodeling, including reduced ciliary coverage and squamous metaplasia.^31–33^ Limited in vivo investigations evaluating the association between PM exposure and inflammation in CRS exhibit greater concentrations of both type 1 and type 2 pro-inflammatory cytokines in nasal secretions during high PM days,^34^ suggesting an “acute on chronic” inflammatory response to ambient PM. At the same time, single cell analyses have demonstrated that at baseline, compared to control tissue, CRS tissue in vivo demonstrates epithelial barrier disruption, tissue remodeling and enhanced inflammation.^35–37^

Nevertheless, whether CRS vs. control tissues respond differently to PM, and how baseline disease dysfunction might influence these responses, is not known. With this in mind, we hypothesized that PM and CRS have additive or synergistic effects on pro-inflammatory cytokine production and epithelial barrier disruption. We incorporated mucociliary-differentiated human sinonasal epithelial cells cultured at an air-liquid interface and employed single cell RNA sequencing (scRNASeq) to study the effects of PM and CRS on the gene expression of specific epithelial subpopulations. Ultimately, our data suggest that CRS tissue is inherently more vulnerable than control tissue to the pathogenic effects of PM.

## Methods

### Patient selection and data collection

We prospectively enrolled adult patients presenting to the University of Michigan (UM) Department of Otolaryngology. Study protocols were approved by UM Institutional Review Board and conducted in accordance with the Declaration of Helsinki. A diagnosis of CRS was confirmed with nasal endoscopy and computed tomography scans, and made in conjunction with criteria outlined by the International Consensus Statement on Allergy and Rhinology,^4^ European Position Paper on Rhinosinusitis and Nasal Polyps,^38^ and the American Academy of Otolaryngology.^39^ After informed consent was obtained, ethmoid sinus tissue from those undergoing endoscopic sinus surgery (ESS) for CRS (i.e., disease) and non-CRS (i.e., control) indications was placed in DMEM with antibiotics and transported to the lab for cell culture at air-liquid interface. Of note, non-CRS indications included surgery for skull base tumors and CSF leaks; control patients had no history of environmental allergies.

### Cell culture

Human nasal epithelial cells were isolated from nasal anterior ethmoids or polyps from controls or patients with CRS. Nasal epithelial cells obtained from nine donors were used for scRNAseq and up to 10 donors used for TEER, ELISA, LDH, or IF. Primary nasal epithelial cells were cultured in Transwells (Corning, Glendale, AZ) at air-liquid interface as described previously.^40^ Briefly, nasal epithelial cells were cultured under submerged conditions in PneumaCult™-Ex Plus medium (Stemcell Technologies, Vancouver, CA) until they reached 90% confluency. Cells were then shifted to PneumaCult™-ALI medium and maintained at air-liquid interface for 21-28 days to achieve fully differentiation, as confirmed by IF staining (Supplemental Figure 1). PM<4 (Standard Reference Material 2786; mean diameter 2.8 µm; National Institute of Standards and Technology) was applied apically in 50 µL Dulbecco’s Phosphate-Buffered Saline (DPBS; final concentration, 30 µg/cm²), with DPBS alone as sham control, for 4 h/day over 3 consecutive days. We decided on our final PM dose by performing a series of initial dosing and exposure experiments. Specifically, we noted that a dose of 30 µg/cm² in differentiated CRS cell cultures generated inflammation, mucus hypersecretion, and epithelial dysfunction, while maintaining minimal cellular cytotoxicity (i.e., <4% lactate dehydrogenase (LDH) assay). Dosing at 10 µg/cm² in these cells showed no upregulation of cytokines. A dose of 30 µg/cm² has also been noted to elicit similar tissue dysfunction in bronchial^41, 42^ and nasal^27^ epithelial cell cultures. Doses greater than 40 μg/cm^2^ have been shown to significantly increase cellular cytotoxicity.^27, 41, 42^ Finally, 30 μg/cm^2^ falls within an expected deposition amount in the human airway for a 12 to 24-hour period in an urban environment.^43^ indeed, in the present investigation a dose of 30 µg/cm² over 72 hours (4 h on/20 h off) induced a pathogenic phenotype while maintaining low cytotoxicity. (Additionally, acute air pollution episodes, such as wildfires, tend to hit their peak ambient PM values at 72 hours,^44^ and this is the timeframe that has been shown to have the strongest association with increased likelihood of emergency department visit for respiratory and cardiovascular events).^44^ Dynasore (Selleckchem) was administered 1 day prior to PM exposure and maintained until sample collection. All experiments were performed in HNECs grown at air-liquid interface until they reached full differentiation after 21-28 days of growth.

### Generation of single-cell suspensions for single-cell RNA-sequencing (scRNA-seq) was performed as follows

HNECs were incubated overnight in 0.4% dispase (Life Technologies) in Hank’s Balanced Saline Solution (Gibco) at 4 °C. Epithelium and submucosa were separated. Epithelium was digested in 0.25% Trypsin-EDTA (Gibco) with 10 U/mL DNase I (Thermo Scientific) for 1 h at 37 °C, quenched with FBS (Atlanta Biologicals), and strained through a 70 μM mesh. Submucosa was minced, digested in 0.2% Collagenase II (Life Technologies) and 0.2% Collagenase V (Sigma) in plain medium for 1.5 h at 37 °C, and strained through a 70 μM mesh. Cells were combined in 1:1 ratio and libraries were constructed by the University of Michigan Advanced Genomics Core on the 10X Chromium system with chemistry v2 and v3. Libraries were then sequenced on the Illumina NovaSeq X to generate 150 bp paired end reads. Data processing including quality control, read alignment (hg38), and gene quantification was conducted using the 10X Cell Ranger software. The samples were then merged into a single expression matrix using the cellranger aggr pipeline.

### Cell clustering and cell type annotation

R package SoupX (v 1.6.2) was used to remove ambient RNA on default parameters. scDblFinder (v 1.16.0) was used to remove doublets using default parameters. The R package Seurat (4.3.0) was used to cluster the cells in the merged matrix. Cells with <500 transcripts or 100 genes, or more than 10% of mitochondrial expression were first filtered out as low-quality cells. The NormalizeData was used to normalize the expression level for each cell with default parameters. The FindVariableFeatures function was used to select variable genes with default parameters. The FindIntegrationAnchors and IntegrateData functions were used to integrate the samples prepared using different 10X Chromium chemistries. The ScaleData function was used to scale and center the counts in the dataset. Principal component analysis (PCA) was performed on the variable genes, and the first 18 PCs were used for cell clustering and uniform manifold approximation and projection (UMAP) dimensional reduction. The clusters were obtained using the FindNeighbors and FindClusters functions with the resolution set to 0.5. Harmony (v 1.0.1) was used to perform batch effect correction, using donor as batch. The cluster marker genes were found using the FindAllMarkers function. The cell types were annotated by overlapping the cluster markers with the canonical cell type signature genes.

### Cluster proportion analysis

To quantify differences in cell composition across conditions, the proportion of each epithelial cell cluster was calculated at the sample (donor) level. For each sample, the number of cells assigned to each cluster was divided by the total number of epithelial cells, and proportions were compared across conditions.

### Gene set enrichment analysis (GSEA)

GSEA was performed to identify biological pathways associated with transcriptional changes between conditions. Differential expression analysis was conducted across individual cells within each cluster to generate ranked gene lists. Gene sets were obtained from the Gene Ontology database (Biological Process). Enrichment analysis was conducted using WebGestalt (v2024). Pathways containing fewer than five genes were excluded. Statistical significance was determined using false discovery rate correction, and pathways with FDR < 0.05 were considered significantly enriched. Normalized enrichment scores were used to compare pathway enrichment across conditions.

### Perturbation score analysis

To estimate the contribution of each epithelial cell cluster to transcriptional differences between conditions, a perturbation score was calculated at the single-cell level for each cluster. DEGs were first identified between groups using the FindMarkers function in Seurat. The perturbation score was then defined as the number of significantly differentially expressed genes divided by the total number of cells within each cluster, allowing comparison of transcriptional perturbation across clusters while accounting for differences in cluster size.

### Pathway activity scoring

To compare pathway activity across conditions and cell types, pathway scores were calculated at the single-cell level using the AddModuleScore function implemented in Seurat. This method computes the average expression of genes within a defined gene set for each cell and subtracts the aggregated expression of matched control gene sets. Module scores were calculated for selected pathways and subsequently averaged across cells within each cluster or condition to enable comparisons of pathway activity.

### Visualization

Dimensionality reduction was performed using uniform manifold approximation and projection to visualize cell clusters. Heatmaps were generated to display the expression of selected gene sets or pathway-associated genes across samples or clusters. Dot plots and bar plots were used to summarize gene expression patterns, pathway scores, and cluster-specific contributions across conditions. All visualizations were generated using the Seurat and ggplot2 packages in R.

### Transepithelial electrical resistance (TEER) assay

TEER was measured 3 days after PM exposure using an EVOM™ Manual meter (Model EVM-MT-03-01; World Precision Instruments, Sarasota, FL, USA) in 12-mm Transwell inserts. After removing the medium, inserts were rinsed three times with DPBS and filled with 500 µL apically and 1 mL basolaterally. The probe was disinfected in 70% ethanol for 10 min and air-dried for 20 min before use. Resistance was recorded with the probe placed vertically in the apical and basolateral chambers, with blank inserts serving as controls.

### Transmission electron microscopy (TEM)

TEM was conducted 3 days after PM exposure. Culture medium was removed and cells rinsed three times with DPBS. Membranes were fixed in 2% paraformaldehyde and 2.5% glutaraldehyde in 0.1 M sodium cacodylate buffer. This was followed by sealing with parafilm and fixing for 2 h at room temperature and then storing at 4 °C prior to transport to the University of Michigan Microscopy Core for TEM grid preparation and imaging using a JEOL JEM-1400Plus LaB□ transmission electron microscope (JEOL Ltd., Tokyo, Japan). PM particles were quantified by counting the total number of particles across 10 grids square (200-mesh/grid) for each donor, and the summed counts were used for comparison between conditions (n=3 donors from each condition).

### Lactate dehydrogenase (LDH) and enzyme-linked immunosorbent assays (ELISA)

The basal media were incubated on the apical side of the inserts at 37°C for 5 minutes. Cytotoxicity was assessed using the LDH Cytotoxicity Detection Kit (Takara, Fort Worth, TX). Cytokine protein concentrations in airway epithelial cell conditioned media were measured by multiplex ELISA (Luminex, Austin, TX). These studies were performed by the University of Michigan Rogel Cancer Center Immunology Core.

### Immunofluorescence staining

Cells were fixed with 100% methanol at −20°C. Paraffin-embedded blocks were sectioned at 500-μm intervals at a thickness of 5 μ, and each section was deparaffinized, hydrated, and stained. Staining with anti-α-acetyl-tubulin (Sigma-Aldrich) and anti-ZO1 (Abcam), anti-KRT5 (Biolegend), and anti-KRT13 (Proteintech) were visualized by immunofluorescence microscopy.

### Data analysis

For PM particle quantification, data are presented as mean ± SD. Statistical significance was assessed using an unpaired two-tailed t-test. For comparisons of epithelial cell cluster proportions, data are presented as mean ± SD. Statistical significance was evaluated using an unpaired two-tailed t-test for comparisons between control and CRS groups, and a paired two-tailed t-test for comparisons between PM-treated and untreated samples from the same donor. For all other experiments, two-way ANOVA was performed to compare the four conditions, with paired comparisons used to assess the effect of PM treatment and unpaired comparisons used to assess the effect of disease status.

## Results

### scRNA-seq analysis of human nasal epithelial cell cultures from CRS and control donors

To characterize epithelial heterogeneity in chronic rhinosinusitis (CRS) vs. controls, and assess responses to PM exposure, we performed scRNAseq on HNEC cultures from nine donors (Figure 1, see Supplemental Table 1 for donor details). After quality control filtering, 344,858 high-quality cells were retained for analysis. Unsupervised clustering identified seven epithelial clusters corresponding to major cell types, including basal, secretory, squamous, cycling basal, deuterosomal and ciliated cells, as well as a rare epithelial population, ionocytes (Supplemental Figure 2A). Cell clusters were annotated based on canonical marker gene expression; the top five marker genes for each cluster are shown in Supplemental Figure 2B and Supplemental Table 2).

**Figure 1.**
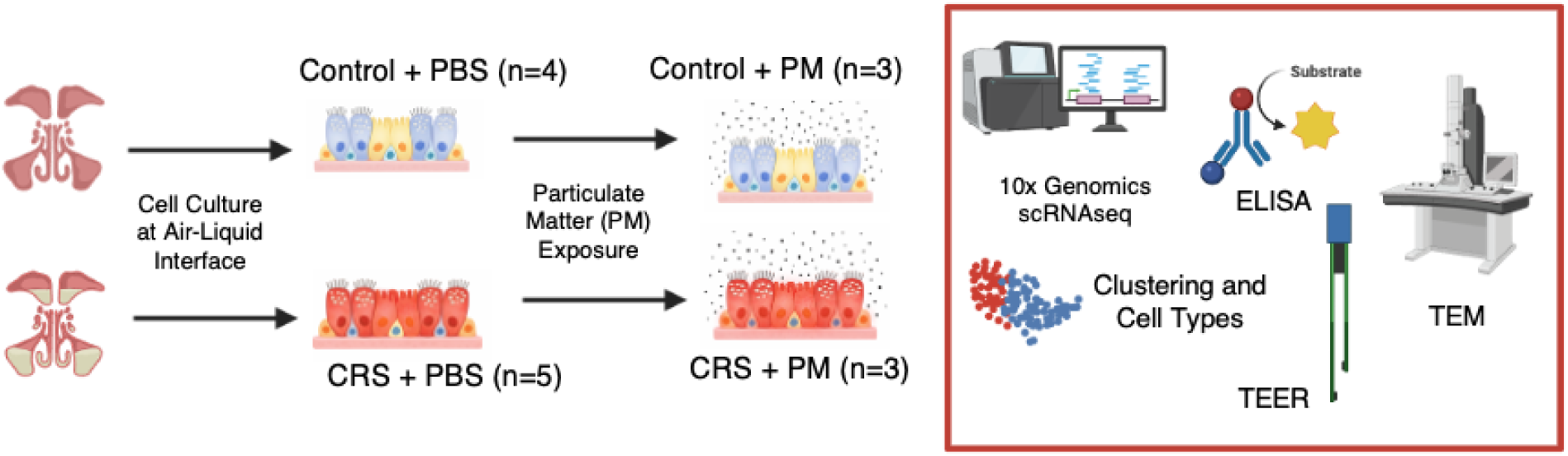
Schematic illustration of the experimental design.

### HNECs from CRS patients exhibit baseline alterations in inflammatory signaling, keratinization, and ciliogenesis

To characterize baseline differences between CRS and control HNECs, we compared baseline (non-PM exposed) cultures from control (n=4) and CRS (n=5) groups (Figure 2A). Squamous cells, a common source of metaplastic change in response to noxious stimuli, were significantly increased in CRS donors, whereas no significant differences were observed in other epithelial clusters (Figure 2B & Supplemental Figure 3A). To determine which cluster contributed most to the baseline expression differences between CRS and control samples, we calculated a perturbation score for each cluster, defined as the number of significantly differentially expressed genes (DEGs) normalized by the number of cells in that cluster. This analysis indicated that basal cells and squamous cells contributed most prominently to the baseline transcriptional differences between CRS and control cultures (Supplemental Figure 4A).

**Figure 2.**
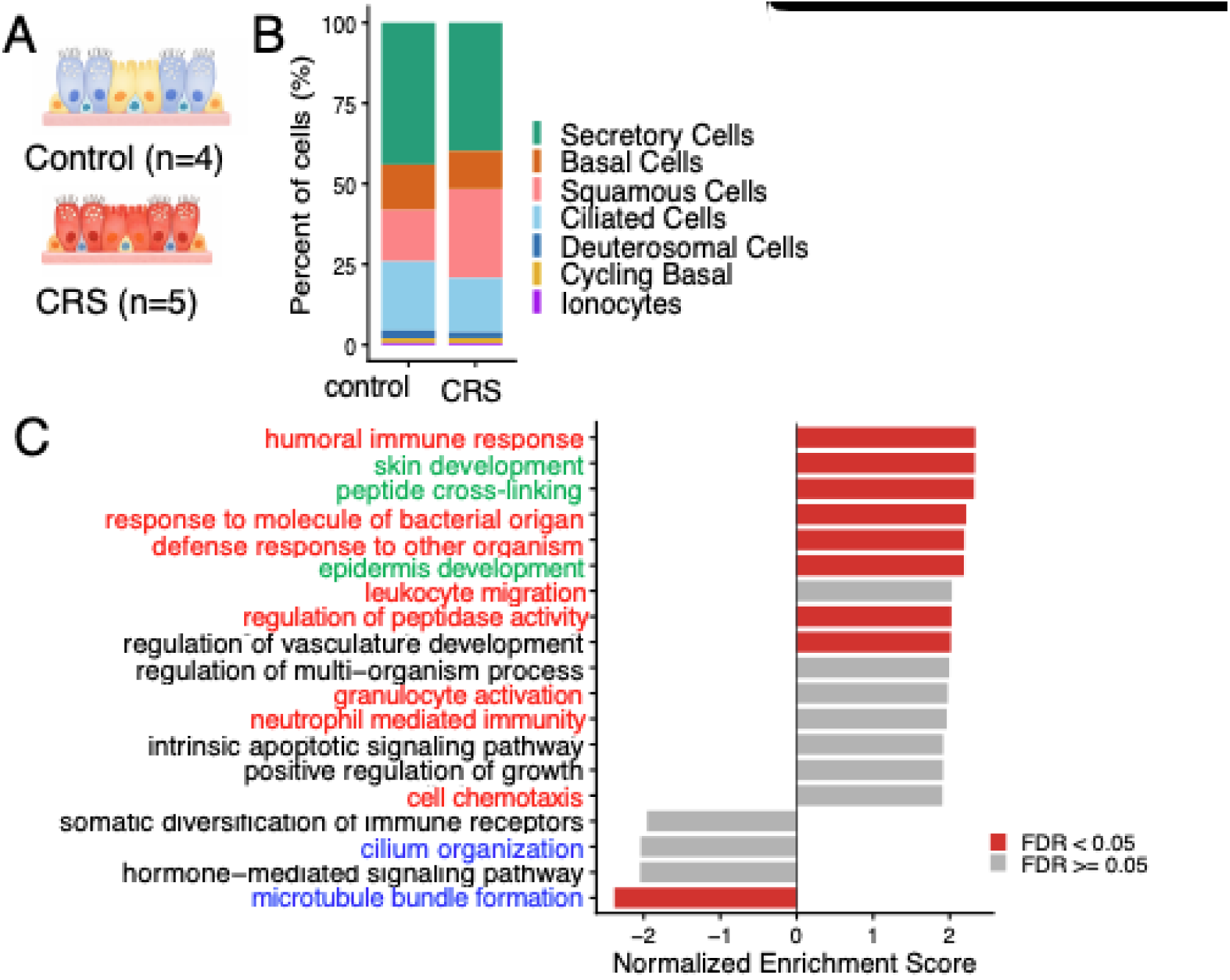
Baseline comparison between control and CRS epithelial cell cultures. A. Schematic illustration of the comparison between control (n=4 donors) and CRS (n=5 donors) cultures. B. Stacking bar plot showing the percentage of each epithelial cell clusters. C. Top enriched pathways identified by GSEA, ranked by normalized enrichment score (NES). Immune signaling pathways are highlighted in red; epithelial remodeling pathways are highlighted in green; cilia-associated pathways are heighted in blue.

We next performed gene set enrichment analysis (GSEA). CRS cells showed increased enrichment of immune-related pathways, including *humoral immune response*, *response to molecule of bacterial origin*, and *defense response to other organism*. In parallel, pathways associated with squamous differentiation and keratinization, such as *skin development* and *epidermis development*, were also increased in CRS cells. Conversely, gene sets related to *microtubule bundle formation* and *cilium organization* were decreased, suggesting impaired ciliogenesis in CRS cultures (Figure 2C).

CRS donors showed increased enrichment of inflammatory pathways related to immune responses, as well as increased squamous metaplasia/keratinization (Supplemental Figure 4B). In contrast, pathways associated with ciliogenesis were decreased in CRS. These data suggest that squamous metaplasia, heightened immune responses and depressed ciliogenesis are consistent features of CRS derived HNECs.

To further determine the cellular sources driving these pathway changes, we compared enrichment scores across epithelial clusters (Supplemental Figure 4C). This analysis revealed that the decrease in genes associated with *microtubule bundle formation* and *cilium organization* resulted primarily from alterations in expression within the ciliated cells, whereas the changes in genes associated with immune responses and keratinization resulted primarily from alterations in the squamous cells.

### PM exposure induces inflammatory signaling and keratinization, while suppressing ciliogenesis in control HNECs

With the understanding that CRS and control HNECs at baseline differ primarily in inflammation, squamous metaplasia/keratinization and ciliary dysfunction, we assessed the transcriptional response of control cells to PM exposure, comparing three control donors treated with and without PM (Figure 3A). No significant changes in clusters were observed, although cycling basal cells showed a trend toward increased abundance following PM exposure (p=0.068) (Figure 3B & Supplemental Figure 3B). Perturbation score analysis indicated that basal cells and deuterosomal cells contributed most prominently to the transcriptional effects induced by PM in control cultures (Supplemental Figure 5A).

**Figure 3.**
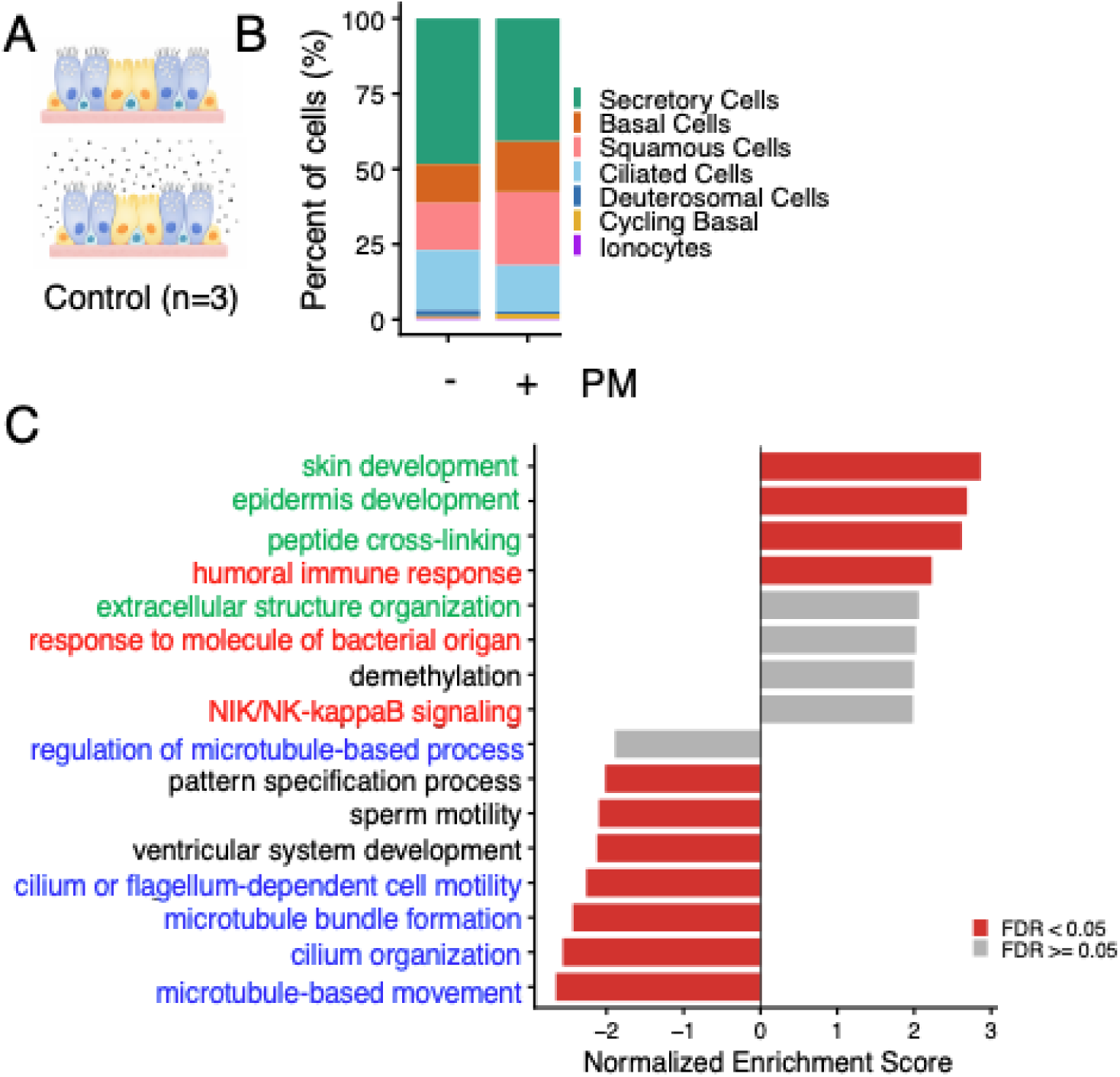
Transcriptional responses of control epithelial cell cultures to PM exposure. A. Schematic illustration of the comparison between PBS– and PM-treated control cultures (n=3 donors). B. Stacking bar plot showing the percentage of each epithelial cell clusters. C. Top enriched pathways identified by GSEA, ranked by NES. Immune signaling pathways are highlighted in red; epithelial remodeling pathways are highlighted in green; cilia-associated pathways are heighted in blue.

GSEA analysis (Figure 3C) revealed that PM exposure also increased enrichment of immune-related pathways, as well as pathways associated with keratinization. In contrast, pathways associated with ciliogeneis were once again decreased. These PM-induced transcriptional changes in controls resembled the baseline transcriptome of CRS cultures. On further subcluster analysis, ciliated cells primarily contributed to the downregulation of pathways associated with cilium organization and microtubule bundle formation, whereas squamous cells contributed most strongly to pathways related to immune responses and keratinization (Supplemental Figure 5B).

### PM exposure induces enhanced IL-1–mediated inflammatory responses in CRS epithelial cells

To assess the transcriptional response of CRS cells to PM exposure, we compared cells from three CRS donors treated with and without PM (Figure 4A). No significant changes in clusters were observed, although basal cells showed a trend toward increased abundance following PM exposure (p=0.052) (Figure 4B & Supplemental Figure 3C). Perturbation score analysis indicated that ciliated cells and squamous cells contributed most prominently to the transcriptional effects induced by PM in control cultures (Supplemental Figure 6A).

**Figure 4.**
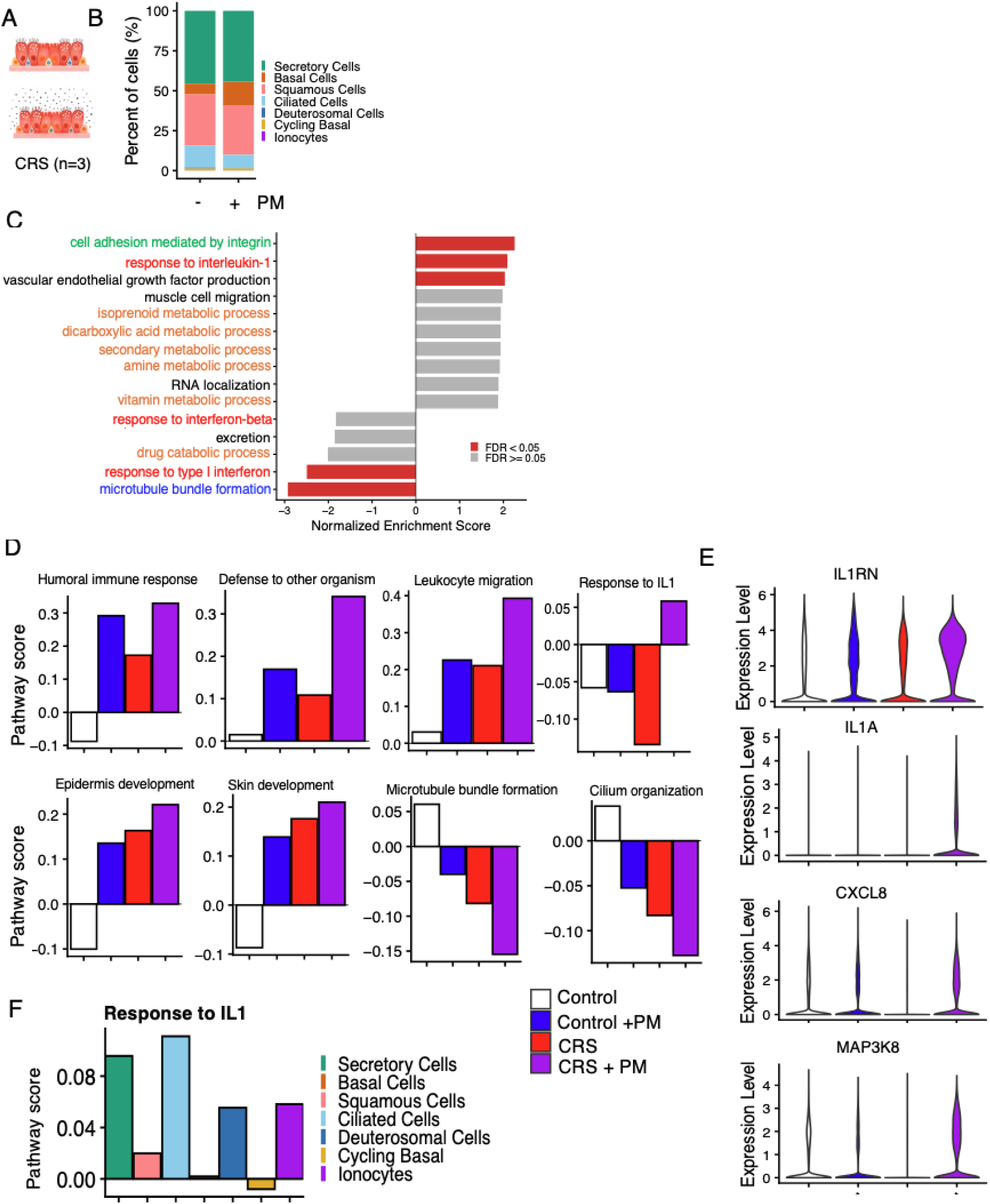
Transcriptional responses of CRS epithelial cell cultures to PM exposure. A. Schematic illustration of the comparison between PBS– and PM-treated CRS cultures (n=3 donors). B. Stacking bar plot showing the percentage of each epithelial cell clusters. C. Top enriched pathways identified by GSEA, ranked by NES. Immune signaling pathways are highlighted in red; epithelial remodeling pathways are highlighted in green; cilia-associated pathways are heighted in blue; metabolism pathways are highlighted in orange. D. Comparison of pathway activity scores across the four conditions: control, PM-treated control, CRS, and PM-treated CRS. E. Expression of represented genes from *response to interleukin 1* pathway across the four groups. F. Comparison of *response to interleukin 1* pathway scores among epithelial cell clusters in PM-treated CRS cultures.

GSEA revealed that PM exposure in CRS cells was also associated with decreased ciliary function and inflammatory signaling upregulation. However, in CRS HNECS the inflammatory pathway specifically and uniquely upregulated was IL-1 (Figure 4D).

We then compared pathway activity scores, calculated as the average expression of genes within each pathway gene set, across the four conditions (i.e., control, control-PM, CRS, and CRS-PM; Figure 4F). CRS cultures exposed to PM exhibited the most pronounced pathway alterations, suggesting a potential baseline vulnerability to the effects of PM compared to controls. Once again, the IL-1 pathway was uniquely increased in this group, as shown by pathway score analysis (Figure 4F) and expression of represented genes (Figure 4G). The pathway activity was predominantly driven by ciliated and secretory epithelial cells in this group (Supplemental Figure 6B).

### CRS cells are more inflamed and sensitive to PM than control

To validate our transcriptional findings at the protein level, cytokine production was measured by ELISA. Two-way ANOVA was used to compare the four experimental conditions, with paired comparisons to assess the effect of PM treatment and unpaired comparisons to assess the effect of disease status (Figure 5A). We first assessed the effect of disease status on cytokine expression without PM. Compared to control tissue, CCL2 was significantly increased in CRS samples. IL-8 was also increased in CRS cells but did not achieve statistical significance. PM exposure significantly increased IL-6, IL-8, CCL-11, CXCL1, IL-1α and IL-1β in CRS epithelial cells but not control cells. Consistent with these findings, IL-6, IL-8, CCL-2, IL-1α, and IL-1β levels were significantly elevated in PM-treated CRS epithelial cultures compared to PM-treated control culture. Together, these data indicate enhanced inflammatory responsiveness in CRS-derived epithelium. Specifically, our transcriptomic and protein data both point to upregulation of the IL-1 pathway after PM exposure.

**Figure 5.**
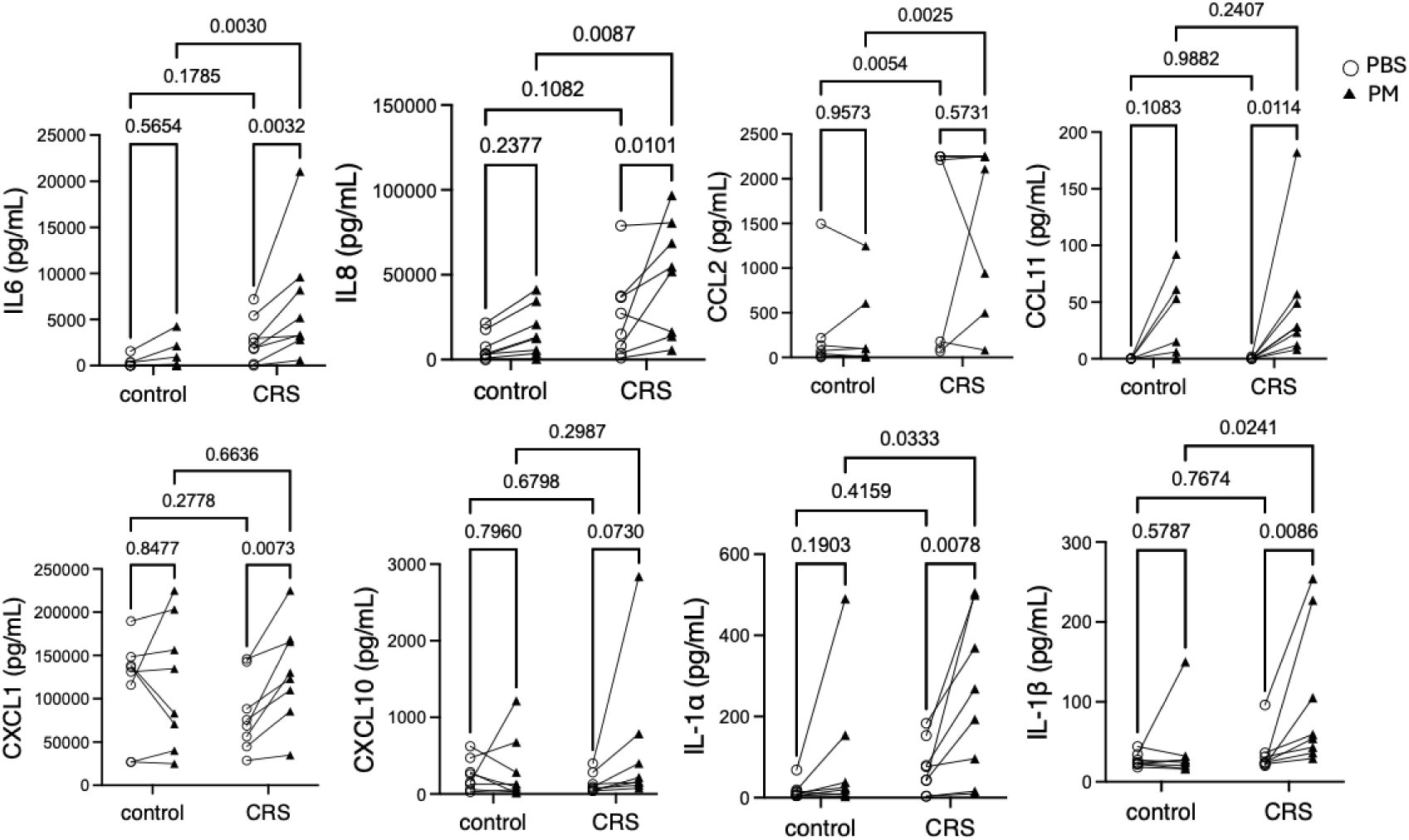
Cytokine assay. ELISA measurements of cytokine levels in culture supernatants from cells treated with or without PM for 3 days (n=10 donors). Statistical significance was assessed by Two-way ANOVA analysis.

To assess epithelial integrity and cellular injury, we performed lactate dehydrogenase (LDH) release assays, transepithelial electrical resistance (TEER), and immunofluorescence (IF) staining. At baseline, no significant differences in cell death were observed between CRS and control cultures. However, PM exposure induced significantly greater cell death in CRS cells compared to controls, indicating increased susceptibility to PM-induced epithelial injury (Figure 6A).

**Figure 6.**
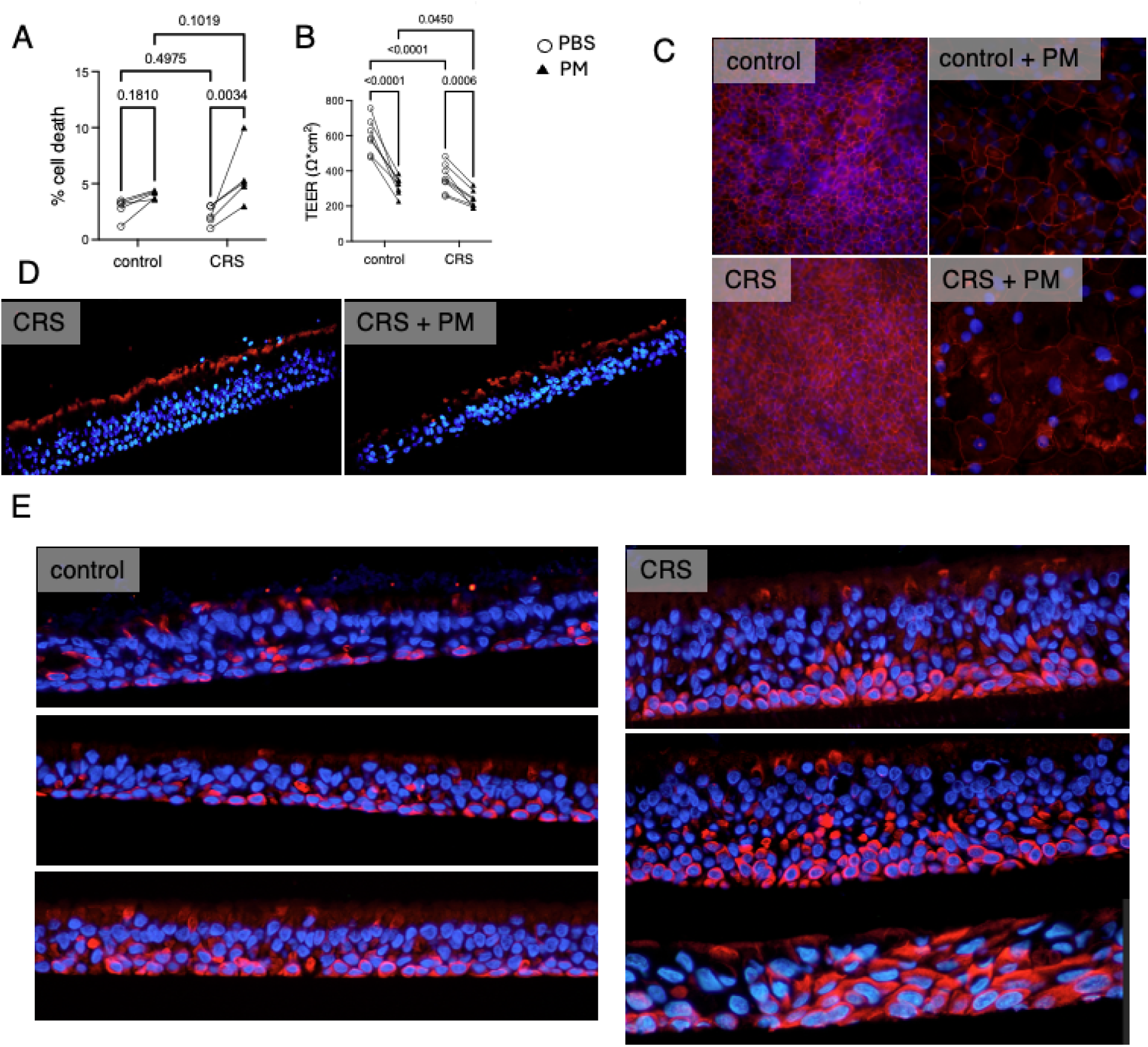
Functional validation of loss of cilia and barrier integrity in epithelial cultures. A. LDH assay showing cytotoxicity in epithelial cultures treated with or without PM for 3 days (control, n=7 donors; CRS, n=8 donors). B. TEER measurements indicating epithelial barrier integrity. Immunofluorescence staining of ZO-1 (red) (C) to assess tight junction structure, acetyl-tubulin (red) (D) to assess cilia loss and KRT13 (red) (E) to assess squamous cell differentiation. Statistical significance was assessed by Two-way ANOVA analysis.

Barrier function was evaluated by TEER measurements. CRS cultures exhibited significantly lower baseline TEER compared with controls, and PM exposure further reduced TEER in both groups, with a further reduction observed in CRS epithelial cultures (Figure 6B). Consistent with these findings, IF staining demonstrated disruption of tight junction protein ZO-1 in both control and CRS epithelial cells, with more pronounced disruption observed in CRS cultures (Figure 6C & Supplemental Figure 2). In addition, staining of acetylated α-tubulin revealed loss of cilia in CRS epithelial cells after PM exposure, suggesting impaired ciliogenesis and epithelial barrier maintenance (Figure 6D). Finally, staining for KRT13 (marker for squamous epithelia) demonstrated an increased squamous state in CRS cultures (Figure 6E).

### Transmission electron microscopy reveals more particulate matter in CRS epithelial cells compared to controls

To better understand the subcellular localization of PM in treated cells, we performed transmission electron microscopy of differentiated HNECs exposed to PM. To do this, we studied PM-treated HNECs cultured at the air-liquid interface from six different donors (three controls and three CRS). Ultrastructural analysis using transmission electron microscopy (TEM) revealed PM particles localized within membrane bound organelles, including vacuoles and lysosome-like structures, suggesting endocytic processing (Figure 7A and Supplemental Figure 3). PM particles were identified based on their morphology and electron-dense appearance (Supplemental Figure 4), consistent with previously characterized features.^45, 46^ Notably, CRS-derived HNECs exhibited a statistically significant (p=0.001) increase in intracellular PM particles compared to non-CRS controls (Figure 7B). To test if changes in TEER were driven by PM internalization, we treated ALI CRS derived HNEC cultures with Dynasore, a small molecule inhibitor of dynamin that halts clathrin-dependent and caveolae-mediated endocytosis. Dynasore significantly restored PM-induced reductions in TEER (Figure 7C) and reduced PM-induced IL-1α and IL8 production (Figure 7D) in CRS donors. Together, these findings suggest that PM internalization may contribute to epithelial barrier dysfunction that is more pronounced in CRS HNECs.

**Figure 7.**
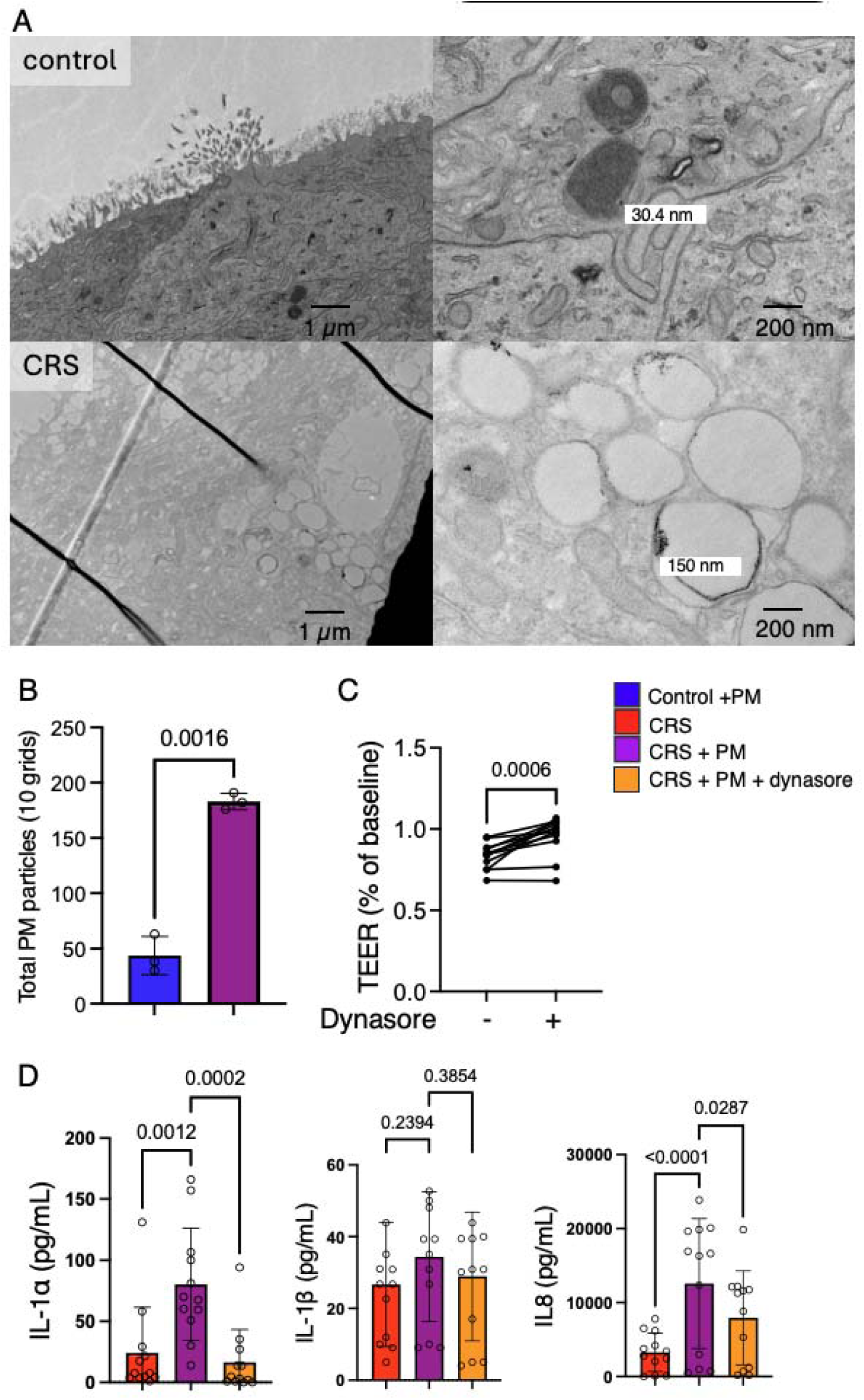
TEM of cultures treated with PM. A. Represented figures showing PM inside cellular cytoplasm. B. quantification of PM. PM particles were quantified by counting the total number of particles across 10 TEM grids (200-mesh) for each donor, and the summed counts were used for comparison between conditions (n=3 donors). C. TEER of PM-treated CRS cultures with or without the endocytosis inhibitor dynasore, measured at day 3 (n=12 donors). D. Protein levels of IL-1α, IL-1β, and IL-8 were quantified by ELISA in PM-treated CRS cultures with or without the endocytosis inhibitor dynasore at day 3 (n = 12 donors). Data were represented as mean ± SD. Statistical significance was assessed by non-paired t test (panel B), paired t test (panel C), or RM one-way ANOVA analysis (panel D).

## Discussion

Although epidemiologic studies^13, 14, 16, 17, 47^ have linked PM exposure with CRS development and disease severity, the underlying cellular mechanisms remain unclear. In this study, using single-cell transcriptomics and complementary functional assays, we demonstrated that CRS-derived HNECs exhibit baseline epithelial dysfunction characterized by altered inflammatory signaling, epithelial differentiation, and impaired barrier integrity. We further showed that PM exposure induced CRS-like molecular and functional features in control epithelial cell cultures, and that, compared to controls, CRS epithelial cells displayed an exaggerated response to PM. Three core pathogenic phenotypes were observed after PM exposure: squamous metaplasia, ciliary dysfunction, and increased inflammatory signaling, culminating in significant epithelial barrier leakiness. We demonstrated that PM localized intracellularly within membrane bound organelles in HNECs, and that rescue of PM-induced epithelial barrier dysfunction and inflammation was possible by pre-treating these HNECs with an endocytosis inhibitor. Ultimately, our findings suggest that CRS HNECs are more vulnerable to PM injury than control HNECs, while highlighting intracellular localization of PM as a potential therapeutic target.

CRS epithelium in vivo is characterized by impaired barrier function, disrupted tight junction architecture, and decreased expression of junctional proteins such as occludin, claudins, and ZO-1, in part mediated by inflammatory cytokines including IL-4 and IFN-γ.^47^ Epithelial cells further contribute to disease pathogenesis through production of cytokines, such as IL-25, IL-33, and TSLP, which promote immune polarization and amplify local inflammation.^51^ Jin et al^48^ used single cell transcriptomics to subcluster epithelial cells in vivo, delineating 8 unique clusters with gene enrichment analysis showing upregulation of cell adhesion and downregulation of actin cytoskeleton and collagen fibril organization in the epithelial population. Similarly, our in vitro cell culture system, comprised of 7 unique epithelial subclusters, demonstrates, at baseline, upregulation of cell adhesion and downregulation of microtubule bundle formation, as well as ciliary loss. These findings highlight the ability of an in vitro HNEC model at air-liquid interface to recapitulate the epithelial nuances of in vivo tissue.

Using this physiologically relevant in vitro system, we examined the effects of PM exposure on epithelial function. We found that PM exposure induced inflammatory signaling, disrupted tight junctions, impaired ciliogenesis and activated keratinization pathways. Previous studies in airway epithelial cells have shown that PM exposure increases inflammatory cytokine production and induces broad transcriptional reprogramming characterized by activation of IL-1–driven inflammatory pathways, mucus secretory programs, and suppression of ciliated cell differentiation.^27,33^ In addition to inflammatory effects, PM has been shown to disrupt epithelial barrier integrity by reducing the expression and localization of tight junction proteins such as ZO-1, JAM-A, and claudin-1.^27,28^ PM exposure can also reduce ciliary coverage, disorganizing cilia structure and downregulating cilia-related genes.^33,34^

Limited prior in vivo work has noted evidence of squamous metaplasia in the sinonasal tissues of subjects living in polluted (i.e., Mexico City) cities compared to control lower polluted environments.^33^ However, to the best of our knowledge, this is the first investigation to demonstrate PM as a driver of airway epithelial plasticity founded in the development of squamous metaplasia, a specific knowledge gap noted in the literature.^49^ This observation highlights another potential contributor to the epithelial barrier dysfunction observed in response to PM exposure, as squamous metaplasia has been shown to decrease epithelial resistance due to epithelial unjamming.^50^

The aforementioned, PM-induced inflammatory, keratinization, and ciliary changes were markedly more pronounced in CRS-derived cultures, suggesting a heightened susceptibility of diseased epithelium to the effects of PM. Single-cell analysis further identified the epithelial subpopulations underlying these changes, with PM-associated effects largely driven by squamous cell populations exhibiting increased inflammatory signaling and altered differentiation, alongside reduced ciliogenesis in ciliated cell populations. Moreover, PM exposure in control epithelial cultures recapitulated key CRS-associated transcriptional and functional features, supporting a potential role for environmental particulate exposure in the initiation of CRS pathogenesis and suggesting that epithelial cells may represent an early and primary site of disease initiation.

To better understand how PM induces epithelial dysfunction, we performed ultrastructural analysis. To the best of our knowledge, we show, for the first time, that PM particles penetrate the mucociliary barrier and are internalized in differentiated epithelial cells, localizing within membrane-bound intracellular compartments. Notably, CRS-derived epithelial cells exhibited a greater intracellular uptake of PM compared with controls. Although no prior studies have demonstrated intracellular presence of PM in CRS derived cells, investigations have noted intracellular localization of PM following exposure in megakaryocytes,^51^ mouse bone marrow–derived macrophages,^52^ and a bronchial epithelial cell line (BEAS-2B cells).^58,59^ In addition, a previous study in submerged BEAS-2B cells showed that inhibition of endocytosis with cytochalasin D attenuated PM2.5-induced IL-8 protein expression. Similarly, pharmacologic inhibition of dynamin partially rescued PM-induced barrier dysfunction, as measured by TEER, as well as IL-1α and IL-8 expression, consistent with the notion that the endocytic pathway may play a role in mediating PM epithelial injury.

Several limitations of our study should be considered. First, while our in vitro air-liquid interface model is useful for studying epithelial-intrinsic responses, it does not recapitulate the immune, stromal and vascular interactions that shape CRS in vivo. Thus, our results may reflect epithelial-level susceptibility rather than whole-disease mechanisms and future in vivo studies are needed to confirm the generalizability of these findings. Second, while our study suggests a role for PM internalization and endocytosis, the mechanistic link between PM uptake and epithelial dysfunction requires further investigation in vivo. Third, given the expense of scRNAseq, our sample size is limited, and larger studies are required to establish the generalizability of our findings across patient and PM-treatment regimes. Fourth, our work assesses acute response to PM and thus cannot comment on changes that would be brought on by chronic exposure, or the extent to which PM drives CRS. Taking these limitations into account, future directions include assessing potential epigenetic mechanisms contributing to baseline dysfunction seen in CRS derived HNECs compared to controls; performing functional analyses to elucidate other pathways that may be impacting PM internalization; and determining how PM disrupt epithelial integrity and stimulate immune responses.

In summary, our study demonstrates that: (1) CRS epithelium exhibits baseline dysfunction that may predispose it to environmental injury; (2) PM exposure both induces CRS-like epithelial changes and exacerbates disease-associated phenotypes; and (3) endocytosis inhibition may rescue PM-induced epithelial barrier dysfunction and inflammation. These findings provide new mechanistic insight into how environmental exposures may contribute to CRS pathogenesis and highlight intracellular PM uptake as a potential therapeutic target for mitigating environmental injury in CRS.

## Declaration of AI and AI-assisted technologies in the writing process

During the preparation of this work, the authors used ChatGPT to assist with language refinement and coding support. After using this tool, the authors reviewed and edited the content as needed and take full responsibility for the content of the publication.

## Abbreviations

CRS: Chronic rhinosinusitis
HNECs: Human nasal epithelial cells
PM: Particulate matter
scRNA-seq: Single-cell RNA sequencing
TEER: Transepithelial electrical resistance
IL: Interleukin
TEM: Transmission electron microscopy

---

**Supplemental Figure 1.** IF staining of HNECs for acetylated-tubulin (red) and MUC5AC (green).

**Supplemental Figure 2.** single cell RNA sequencing analysis of human nasal epithelial cell cultures from CRS. A. UMAP visualization of epithelial cell clusters of from nasal epithelial cultures (n=15 samples). B. Heatmap showing the expression of the top five marker gene for each cluster.

**Supplemental Figure 3.** Percentage distribution of epithelial cell clusters. A. Comparison between control and CRS cultures. B. Comparison between PBS– and PM-treated control cultures. C. Comparison between PBS– and PM-treated CRS cultures. Data are presented as mean ± SD. Statistical significance was assessed using an unpaired two-tailed t-test for panel A and paired two-tailed t-tests for panels B and C.

**Supplemental Figure 4.** Baseline comparison between control and CRS epithelial cell cultures. A. Comparison of perturbation scores among clusters, visualized on UMAP (left) and as a bar graph (right). B. Heatmap showing the average expression of genes within selected GSEA pathways. Each column represents an individual donor and each graph represents a pathway.

**Supplemental Figure 5.** Transcriptional responses of control epithelial cell cultures to PM exposure. A. Comparison of perturbation scores among clusters, visualized on UMAP (left) and as a bar graph (right). B. Comparison of enrichment score for selected GSEA pathways across epithelial clusters.

**Supplemental Figure 6.** Transcriptional responses of CRS epithelial cell cultures to PM exposure. A. Comparison of perturbation scores among clusters, visualized on UMAP (left) and as a bar graph (right). B. Comparison of enrichment score for selected GSEA pathways across epithelial clusters.

**Supplemental Figure 7.** ZO-1 staining of cell culture from additional donors.

**Supplemental Figure 8.** TEM of cell culture from additional control donors.

**Supplemental Figure 9.** TEM of cell culture from additional CRS donors.

**Supplemental Figure 10.** TEM of CRS cell culture treated with PM.

**Supplemental Table 1.** Patient demographic information.

**Supplemental Table 2.** Marker genes associated with each cell cluster.

