## Supplemental Figures for "Disease-specific differences in particulate matter handling drive pathogenic responses in human derived nasal epithelial cells"

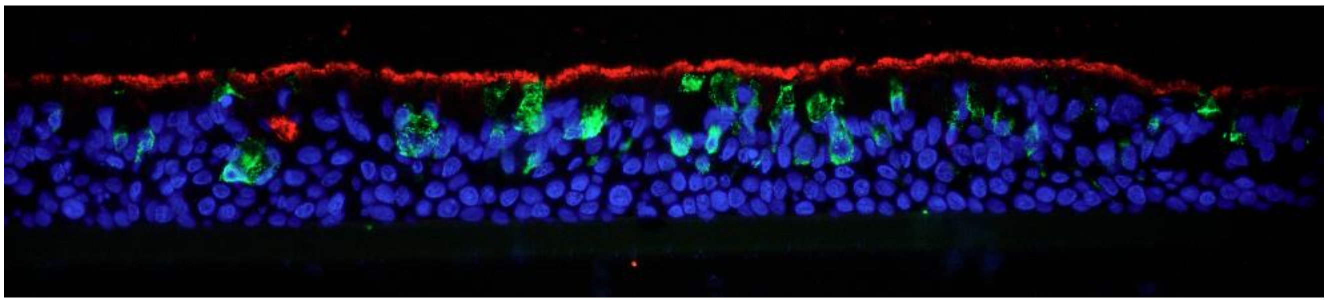

Supplemental Figure 1

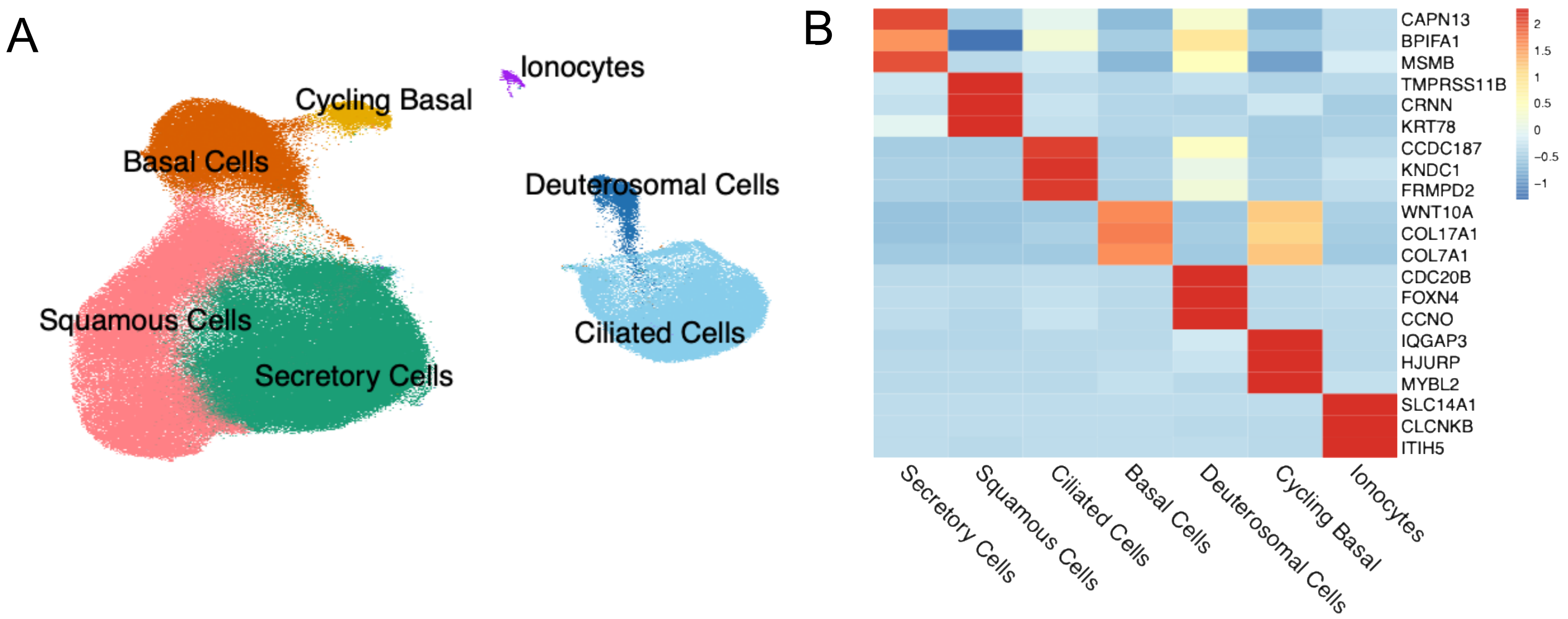


Supplemental Figure 2

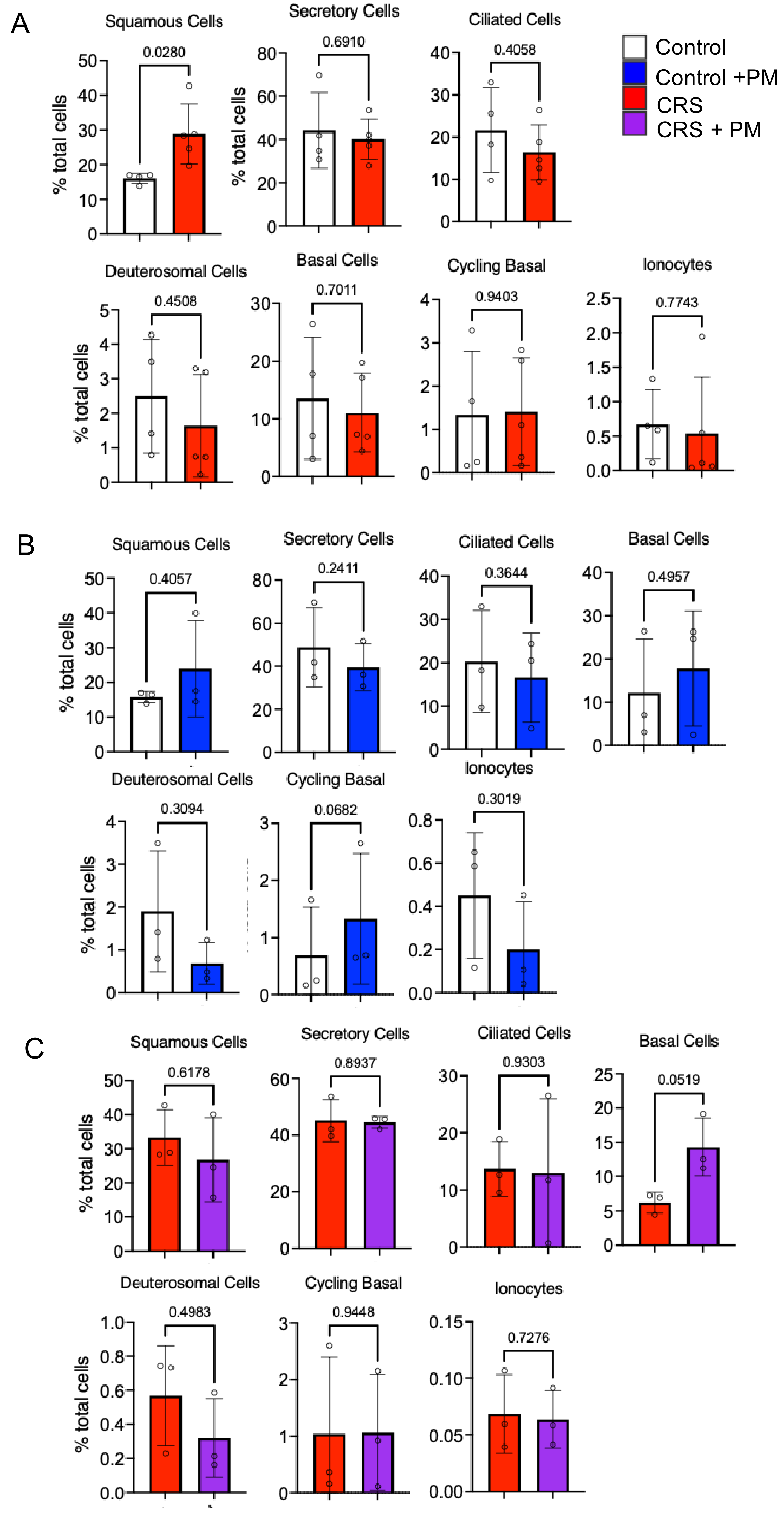


Supplemental Figure 3

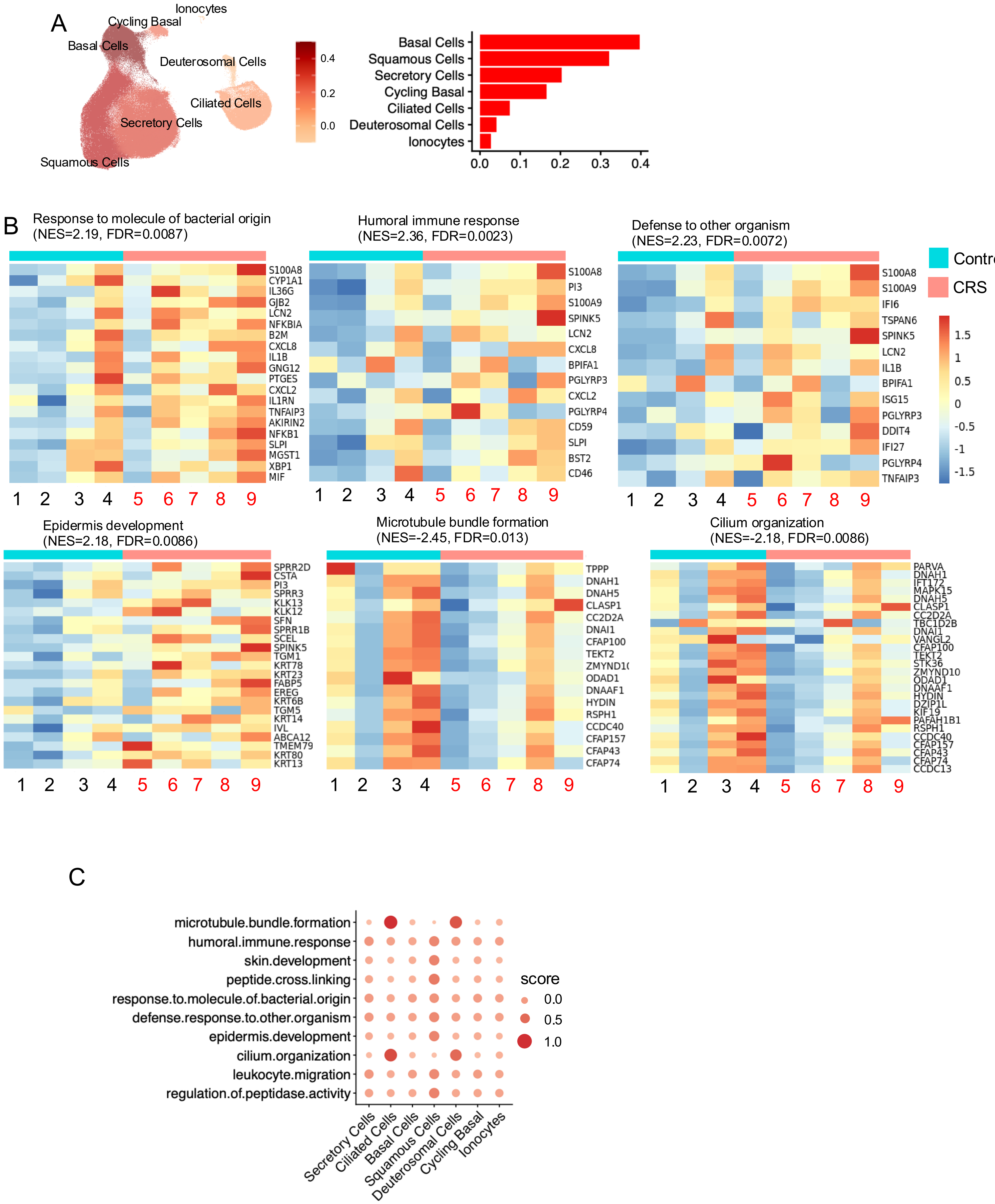


Supplemental Figure 4

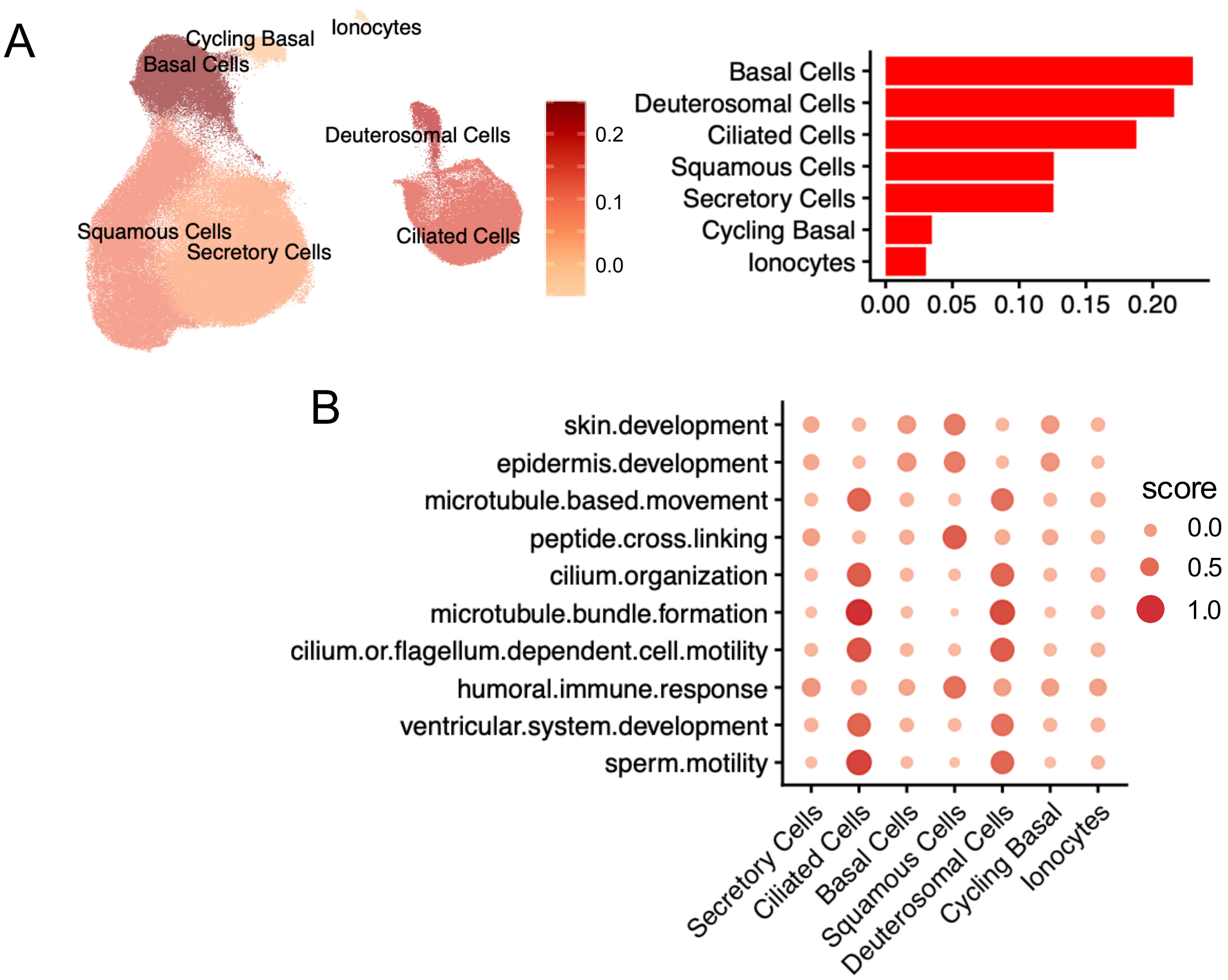


Supplemental Figure 5

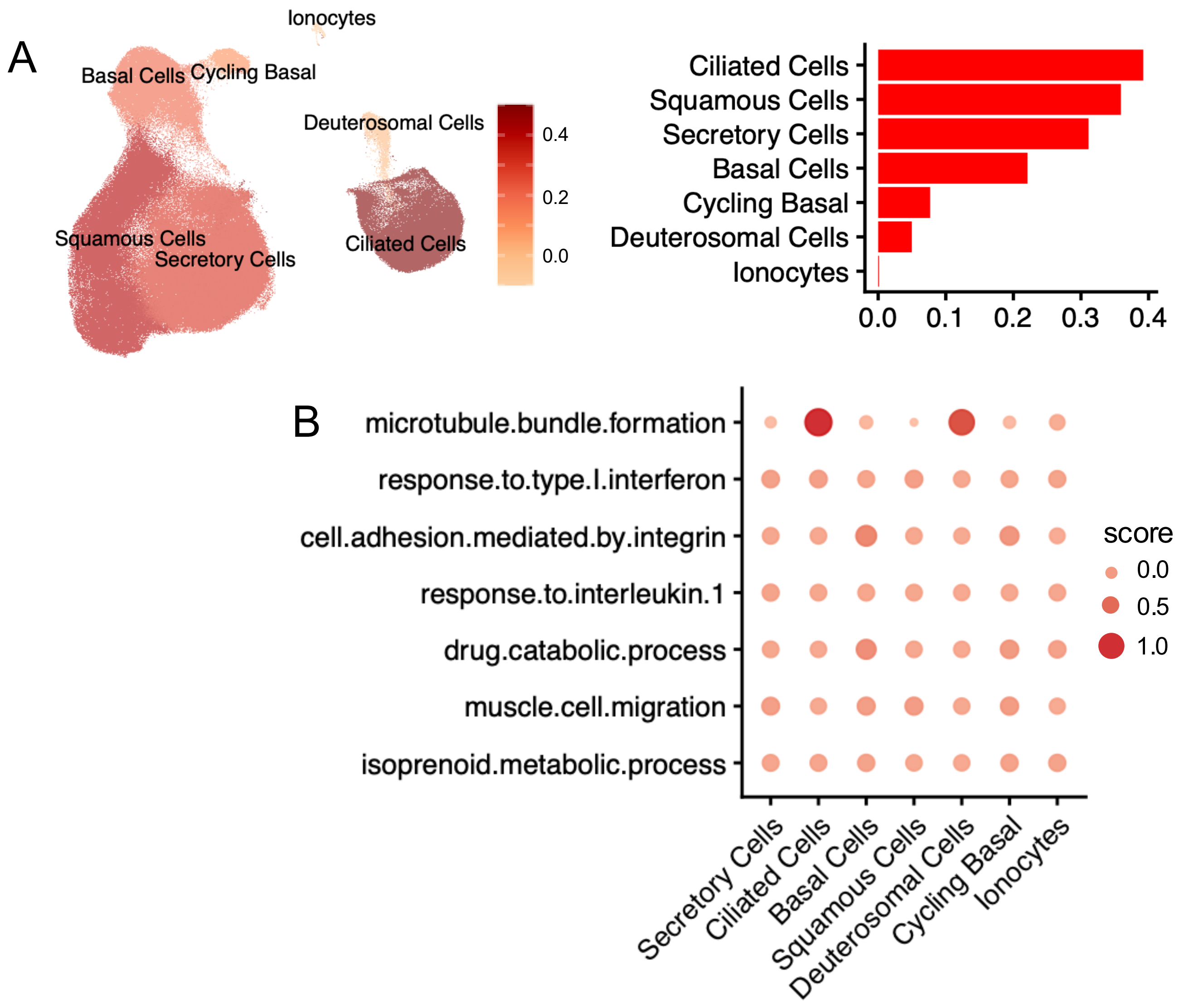


Supplemental Figure 6

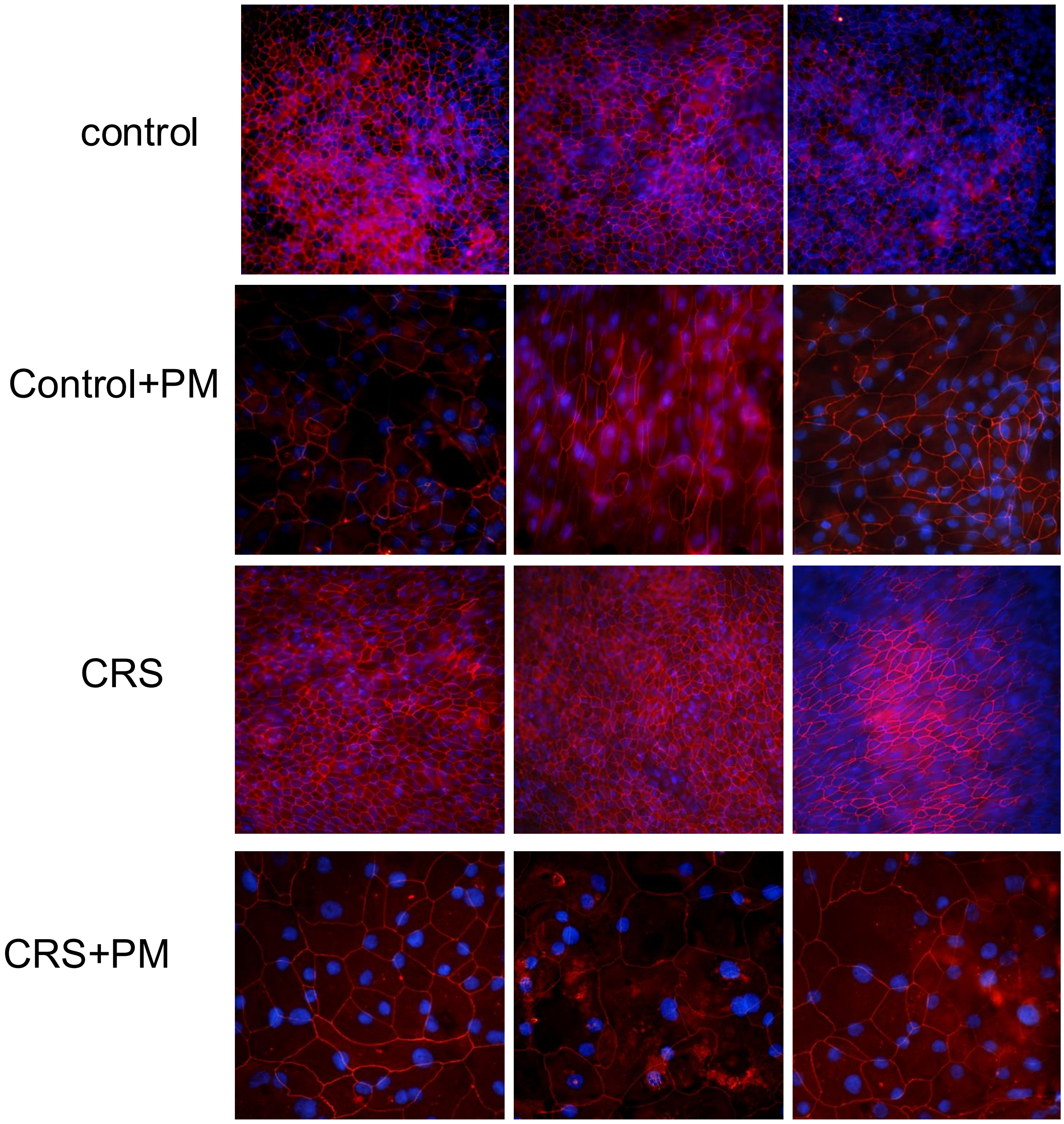


Supplemental Figure 7

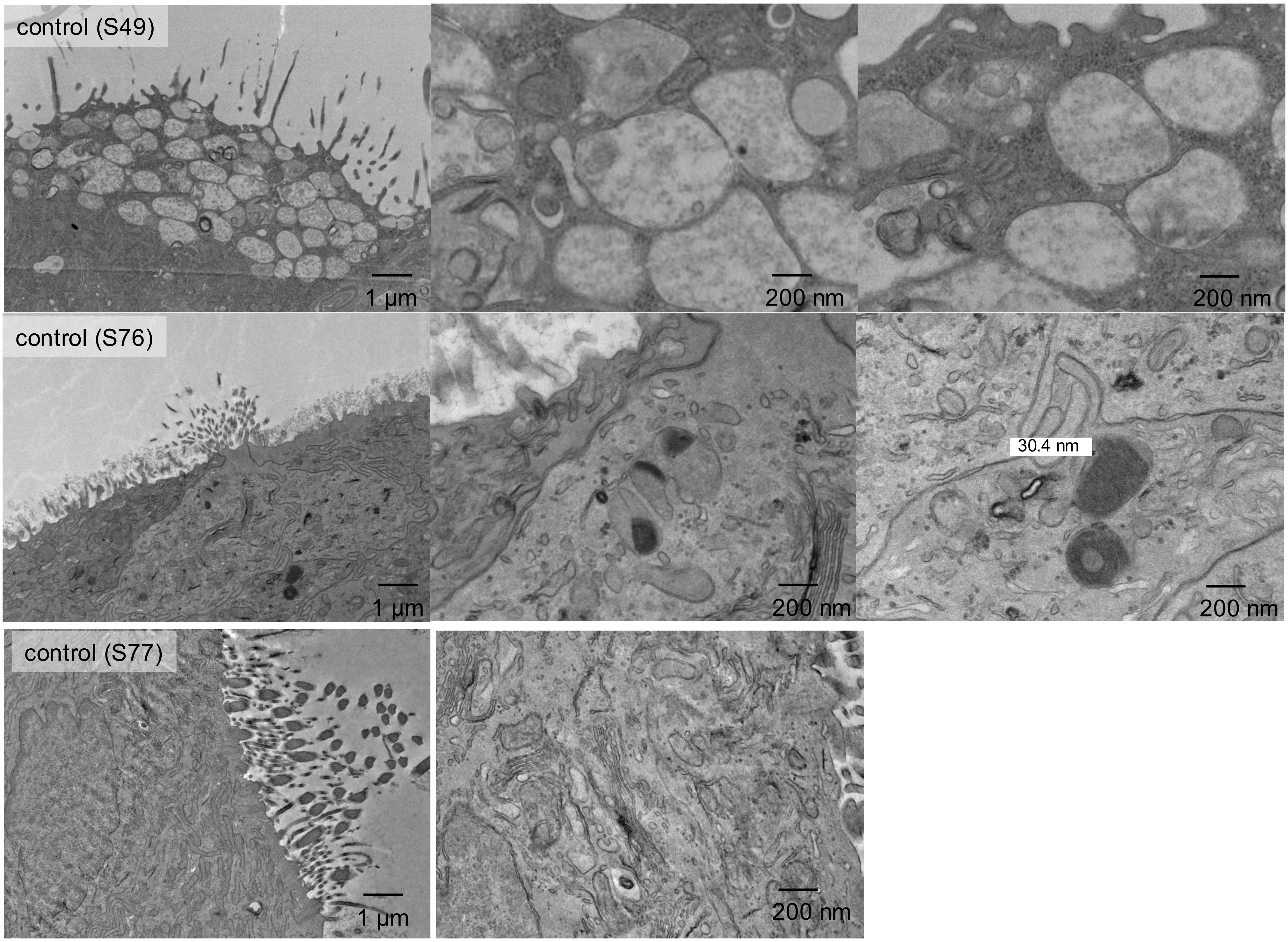


Supplemental Figure 8

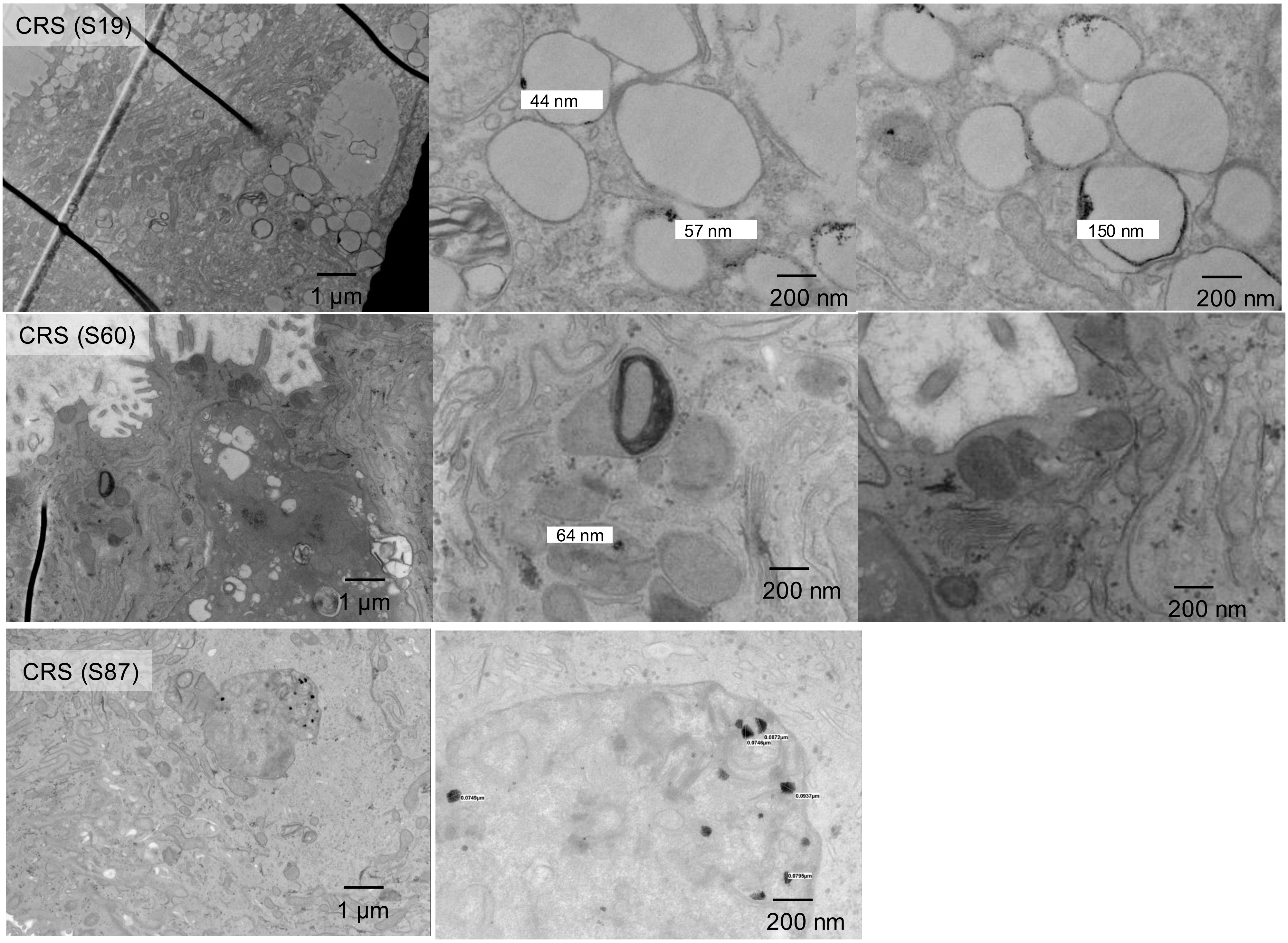


Supplemental Figure 9

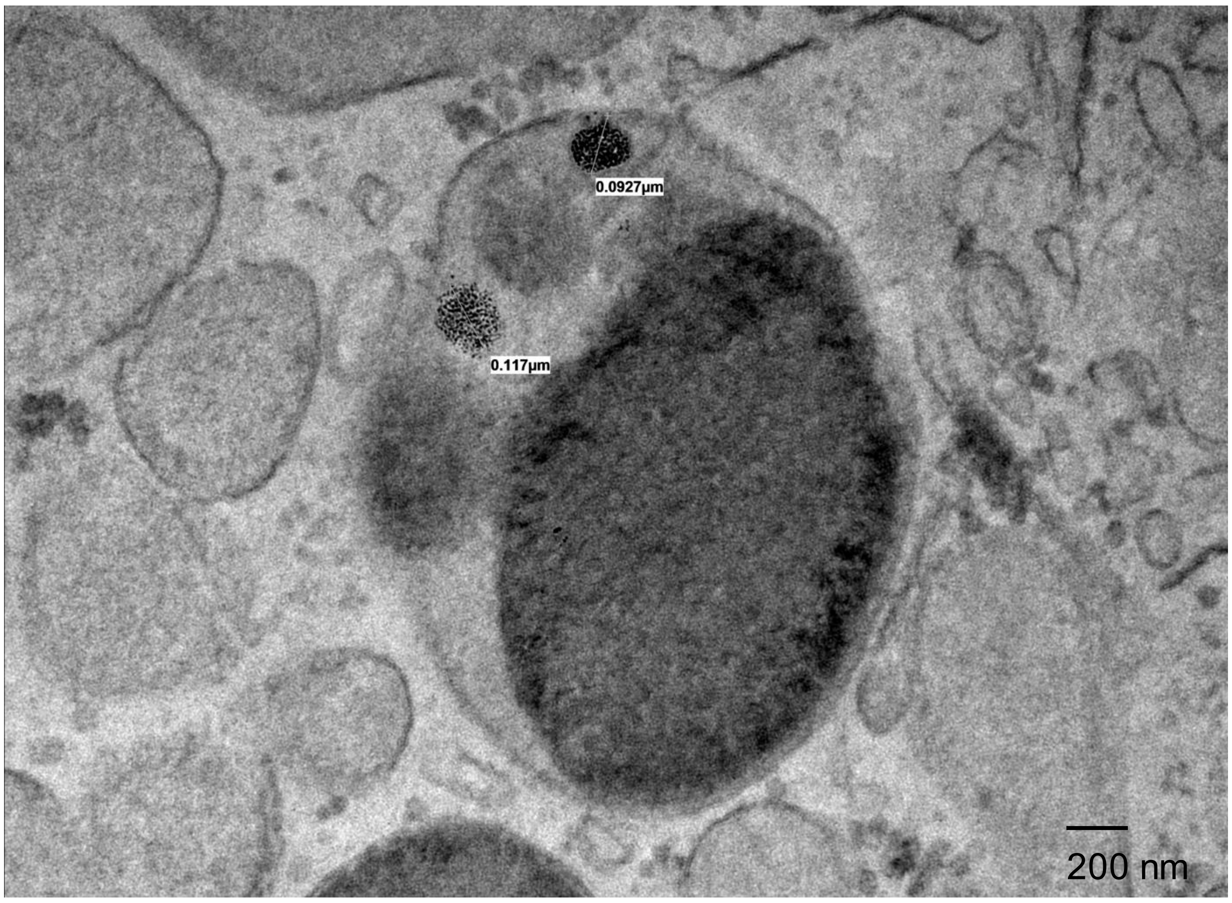


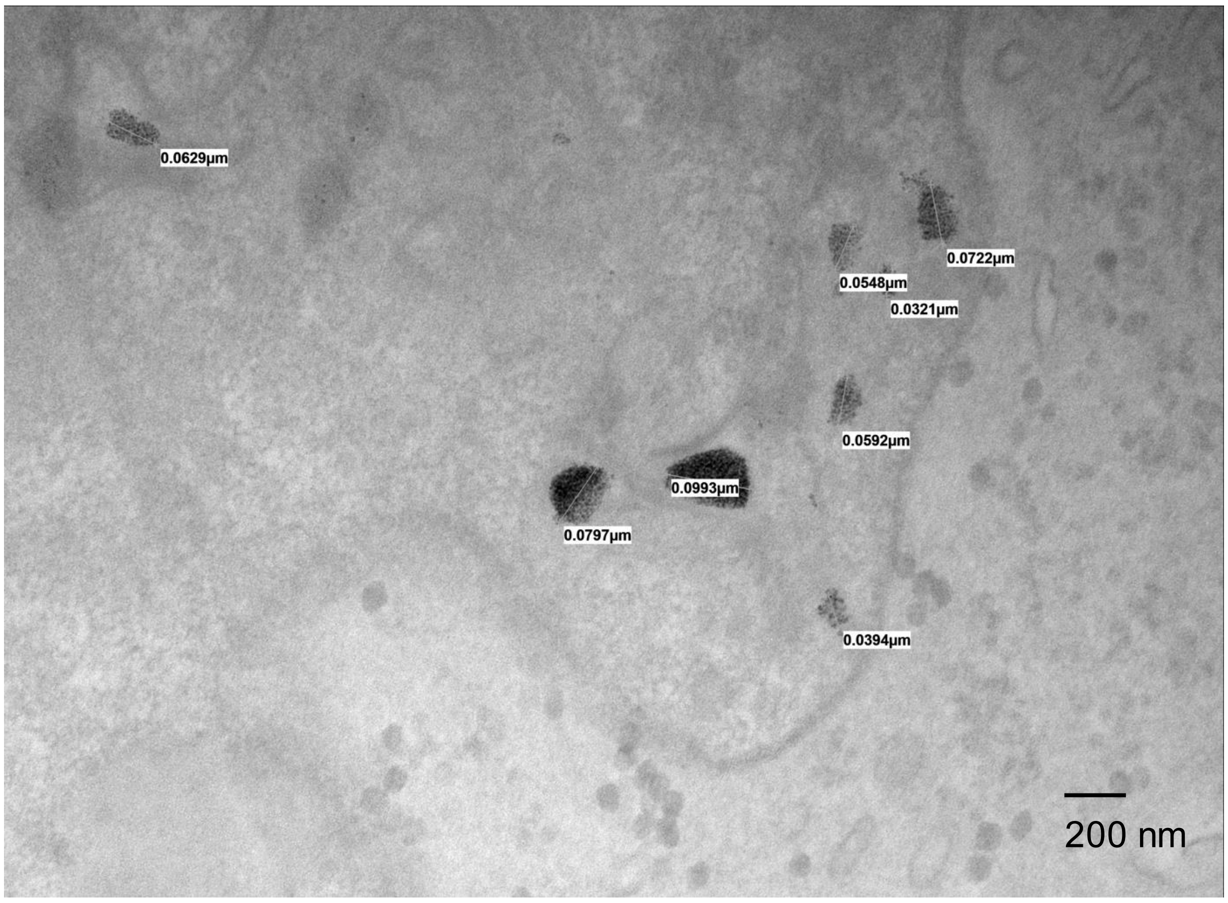


Supplemental Figure 10
